# ADP heptose, a novel pathogen-associated molecular pattern associated with *Helicobacter pylori* type 4 secretion

**DOI:** 10.1101/405951

**Authors:** Lennart Pfannkuch, Robert Hurwitz, Jan Traulsen, Paul Kosma, Monika Schmid, Thomas F. Meyer

**Affiliations:** Department of Molecular Biology, Max Planck Institute for Infection Biology, Charitéplatz 1, 10117 Berlin, Germany; Department of Infectious Diseases and Pulmonary Medicine, Charité, University Hospital Berlin, 13353 Berlin, Germany; Berlin Institute of Health, 10117 Berlin, Germany; Department of Chemistry, University of Natural Resources and Life Sciences-Vienna, 1190 Vienna, Austria

**Keywords:** NF-κB, ALPK1, TIFA, gram-negative, LPS

## Abstract

The gastric pathogen *Helicobacter pylori* activates the NF-κB pathway in human epithelial cells via the α-kinase 1 (Alpk1)–TIFA axis. We and others have previously shown that heptose 1,7-bisphosphate (HBP) acts as a pathogen-associated molecular pattern (PAMP). HBP is an intermediate of lipopolysaccharide (LPS) synthesis in *H. pylori* and other gram-negative bacteria. Deletion of the *hldE* (*rfaE*) gene encoding the enzyme responsible for HBP synthesis, as well as deletion of further upstream genes, causes loss of NF-κB stimulation by *H. pylori*, while deletion of the downstream phosphatase encoding gene *gmhB* does not. This has led to the conclusion that HBP is the PAMP responsible for NF-κB induction. Here, our attempts to identify HBP in lysates of *H. pylori* revealed surprisingly low amounts that fail to explain NF-κB activation. Instead, we identified ADP heptose, a major downstream metabolite of HdlE, as the predominant PAMP in *H. pylori* lysates, exhibiting ∼100-fold stronger activity compared to HBP. It therefore appears that synthesis of ADP heptose from HBP in *H. pylori* occurs independently of GmhB. The data lead us to conclude that ADP heptose constitutes the key PAMP, secreted via the pathogen’s cagPAI encoded type 4 secretion (T4SS).

## Introduction

The innate immune system provides mammalian cells with various means of self-defense against microbial infections. A key element in detecting possible threats is recognition of pathogen-associated molecular patterns (PAMPs) by host cells. These highly conserved structures are found in bacteria, viruses and fungi and can be detected via pattern recognition receptors (PRRs) (Medzhitov, 2007). While a variety of PAMPs have been identified in the last decades, such as lipopolysaccharide (LPS), CpG nucleotides, or flagellin the picture is still incomplete (Baccala et al., 2009). Recognition of these structures alerts the host cell to the presence of potentially harmful microorganisms and triggers innate immune signaling pathways that induce production of inflammatory mediators and attract immune cells to fight an infection.

One of the main pathways activated in response to infection with pathogenic bacteria is nuclear factor κB (NF-κB) signaling, which has been suggested to play a role in inflammation and carcinogenesis (DiDonato et al., 2012). It comprises three pathways: the canonical, non-canonical and p105 pathway all of which are important for the response to infections and the induction of inflammatory mediators but may also be relevant for the modulation of the inflammatory response (Banerjee et al., 2014; O’Reilly et al., 2018). Canonical NF-κB signaling can be activated by a variety of effectors, such as LPS or tumor necrosis factor α (TNF-α) (Newton and Dixit, 2012). Activation of the upstream receptors leads to phosphorylation of inhibitor of NF-κB kinase β (IKKβ) and subsequent phosphorylation of inhibitor of NF-κB α (IκBα). This induces polyubiquitination and proteasomal degradation of IκBα, which releases the p65/p50 heterodimer normally sequestered in the cytoplasm. Upon release, p65/p50 is translocated into the nucleus where it activates the expression of multiple genes involved in inflammation and proliferation (Jost and Ruland, 2007).

LPS is a well-studied PAMP found in most gram-negative bacteria. It has a tripartite structure consisting of lipid A, an oligosaccharide core comprising an inner and an outer core region, and the O antigen. Lipid A is the hydrophobic component of LPS localized in the outer bacterial membrane while core polysaccharides and O antigen polysaccharide are presented on the bacterial surface (Wang and Quinn, 2010). The inner core typically consists of 3-deoxy-D*-manno-*octulosonic acid and L,D-heptose units (Kneidinger et al., 2002).

The precursor of the L,D-heptose units, ADP-L-*glycero*-β-D-*manno*-heptose (ADP-heptose), is synthesized in a five-step pathway, starting from D-sedoheptulose 7-phosphate which is (i) converted into D-α,β-D-heptose 7-phosphate by the isomerase GmhA, (ii) phosphorylated to form D-*glycero*-D-*manno*-heptose 1,7-bisphosphate (HBP) by the bifunctional kinase/adenylyltransferase HldE, (iii) dephosphorylated to D-β-D-heptose 1-phosphate by the phosphatase GmhB, followed by (iv) nucleotide activation by formation of ADP-D-β-D-heptose by HldE and finally (v) epimerization to ADP-L-β-D-heptose by the epimerase HldD (Kneidinger et al., 2002) (Supplementary Figure 1A). The resulting activated heptose unit is then integrated into the LPS core region.

In 2015, the intermediate HBP was identified as a potent effector of NF-κB activation in epithelial cells upon infection with *Neisseria spp.* (Gaudet et al., 2015). More recently, we showed that the proteins α-kinase 1 (ALPK1) and TRAF-interacting protein with forkhead associated domain (TIFA) are key components in the response pathway to HBP and in activating canonical NF-κB signaling in gastric epithelial cells infected with *H. pylori* (Zimmermann et al., 2017). Moreover, activation of NF-κB was found to depend on cytosolic detection of HBP. In *H. pylori* infection, the type IV secretion system (T4SS) is needed for translocation of HBP into host cells (Zimmermann et al., 2017). In *Neisseria* infection, cytosolic delivery via phagocytosis followed by lysosomal degradation as well as endocytosis of HBP has been proposed (Gaudet et al., 2015).

Here we biochemically analyzed the amounts of HBP in lysates of *H. pylori*. Surprisingly, we observed only minor amounts of HBP, which hardly explained the strong inflammatory response of host epithelial cells. We thus hypothesized that yet another compound involved in LPS synthesis might account for the innate immune response observed in *H. pylori* infection. Further analysis revealed abundant ADP-heptose in the lysate. Here, we demonstrate that ADP-heptose is a novel, potent NF-κB-activating PAMP that is recognized by the ALPK1-TIFA signaling axis. Since ADP-heptose is present in all gram-negative bacteria, this molecule might be of major relevance in other gram-negative infections.

## Results

### Identification of ADP heptose as novel NF-κB activator

In order to confirm the presence of HBP in *H. pylori* lysates, we tried to separate the heptose by solid phase extraction (SPE) methods based on the protocol described by Gaudet et al. (2015). Detection of HBP was performed by ion-pairing reversed phase chromatography via electrospray ionization mass spectrometry (ESI-MS). However, in SPE-extracts derived from at least 1 ml of *H. pylori* lysate we could not detect HBP as judged by the appearance of a mass with m/z 369 Da [M-H]^-^ (data not shown). Since the detection limit of HBP was relatively high (≥100 pM), we aimed to increase the sensitivity by derivatizing the heptose by reductive amination using 3-amino-9-ethylcarbazole (AEC) after treatment with either alkaline phosphatase (CIP), which cleaves both the 7’-O-phosphate and the glycosidic phosphate ester, or mild acid, which hydrolyzes only the glycosidic one (Han et al., 2013). This creates an aldehyde group enabling Schiff’s base generation between carbohydrates and primary amines, a reaction which does not occur if the sugar is esterified to phosphate by an *O-*glycosidic bond. We then determined the sensitivity of our assay using chemically synthesized β-HBP as a reference. Derivatization of β-HBP allowed us to detect the AEC-sugar derivates by LC-MSMS (MALDI) in the fM range. We detected precursor masses of m/z 405 [M+H]^+^ for the CIP-treated HBP, with both phosphates cleaved and m/z 485 [M+H]^+^ after trifluoracetic acid (TFA) hydrolysis, removing only the *O-*glycosidic bound phosphate. Reductive amination of the *H. pylori* SPE-extract after acid hydrolysis led to the appearance of the predicted m/z 485 mass and fragment ions (m/z 210, 223,275 and 315) (Fig. 1A) but it also revealed that HBP was present in quantities too low to account for the stimulation of NF-κB. In our experiments with synthetic HBP, a concentration of at least 10 µM was required (Zimmermann et al., 2017). By contrast, in extracts HBP was only detected at concentrations in the fM range. To our surprise, in the lysates we detected a mass of m/z 405 Da (fragment ions of m/z 210, 223, 315) after AEC-derivatization, which appeared after acid treatment (Fig. 1B), indicative of a heptose with all phosphate residues being acid labile.

**Figure 1:**
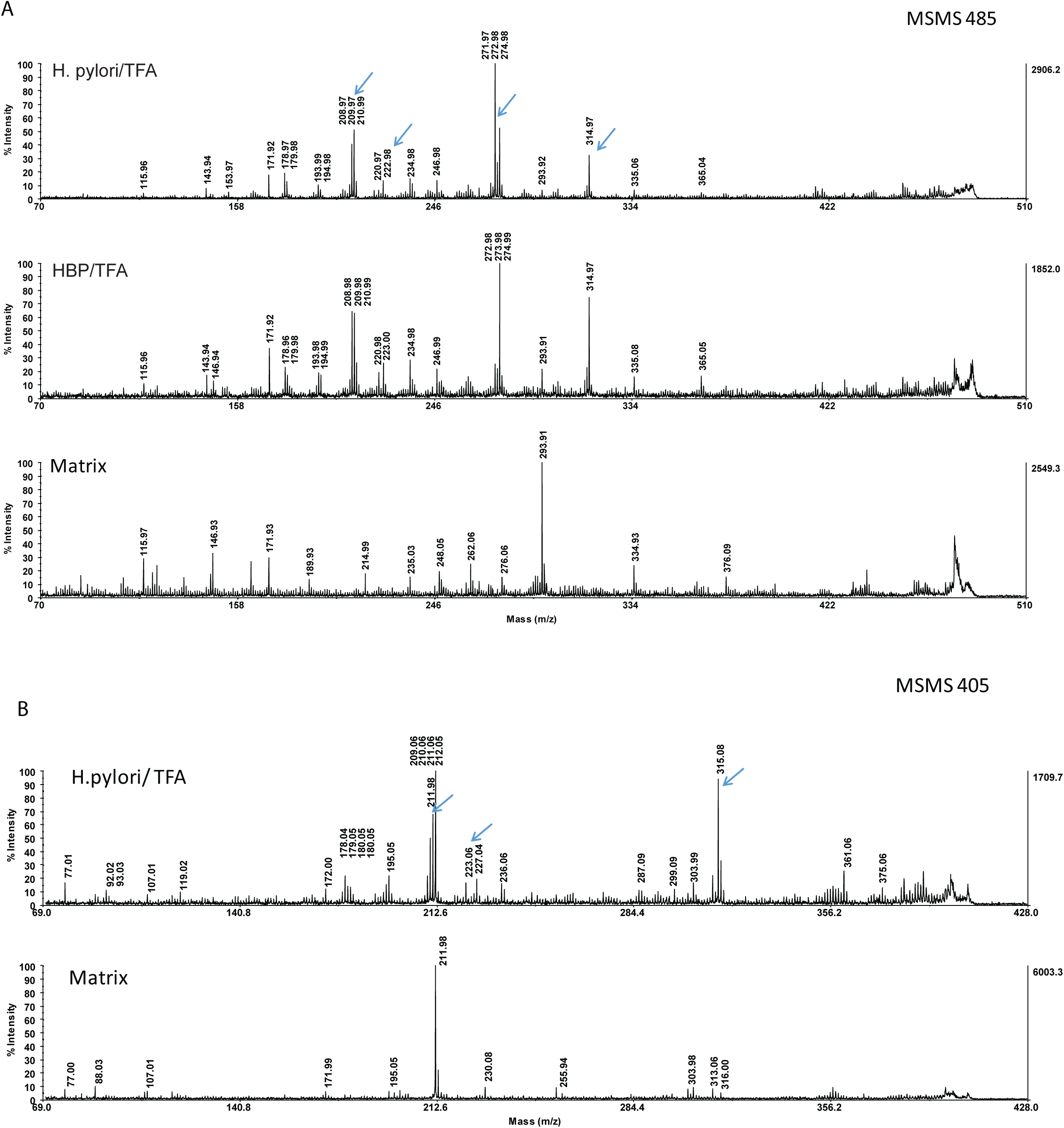
Evidence for HBP in *H. pylor*i extracts, MALDI-TOF MSMS of AEC derivatized heptoses. (A) HBP or extracts from *H. pylori* (graphite carbon eluate) were derivatized with AEC after acid hydrolysis, respectively. AEC-heptose adducts were separated with UPLC and analyzed by MALDI-TOF MSMS. Shown are the MSMS spectra of the precursor ion m/z 485 [M+H]^+^, as the putative Schiff base-derived H7P-AEC adduct. Upper panel: Derivatization of *H. pylori* extracts, blue arrows: Fragment ions (m/z 210, 223,275 and 315) of the precursor ion m/z 485 [M+H], middle: Derivatization of HBP, lower: Matrix control.(B) Upper panel: MALDI-MSMS spectrum of the AEC-*glycero*-D-*manno*-heptose precursor ion (m/z 405 [M+H]^+^) derived from reductive amination of a *H. pylori* extract after acid hydrolysis with trifluoroacetic acid (TFA), blue arrows: Fragment ions (m/z 210, 223, 315) of the precursor ion m/z 405 [M+H]. Lower: Matrix control.

Improved chromatographic resolution of the *H. pylori* carbohydrate compounds from the SPE-extract was achieved by ion-pairing reversed phase-UPLC. We tested the fractions for their ability to stimulate NF-κB in luciferase reporter cells and for the appearance of AEC-derivatized heptoses by MALDI mass spectrometry. By transfecting the fractions into reporter cells we found that only a single faction in any given run had a strong capacity for inducing NF-κB (Fig. 2 A). For these NF-κB stimulating fractions, the most prominent MSMS-peak intensities (m/z 223 Da) of the AEC-precursor masses was quantified (summarized in Supplementary Table 1; see also Supplementary Figure 1 B and C and 2). From these data we concluded that not HBP but rather a *manno*-heptose with an acid labile glycosidic linkage that is partly resistant to alkaline phosphatase was present in the NF-κB-stimulating fractions. This was further supported by the observation that the active fraction contained a prominent compound with a mass of m/z 618 [M-H]^-^ (ESI-MS) and a UV-absorbance of 259 nm, whereas spiked HBP eluted earlier (Figures 2B and C). We therefore hypothesized that the substance could be β-ADP-heptose, the final metabolite of the ADP-heptose pathway. This assumption was further confirmed by the fact that treatment of the active *H. pylori* fraction after UPLC chromatography with alkaline phosphatase (CIP) or with phosphodiesterase from *Crotalus adamanteus* (PDE) showed that PDE completely hydrolyzed the active compound (Supplementary Figure 2 C). By contrast, CIP-treatment caused only minor hydrolysis after prolonged digestion times (Supplementary Figure 2 C).

**Figure 2:**
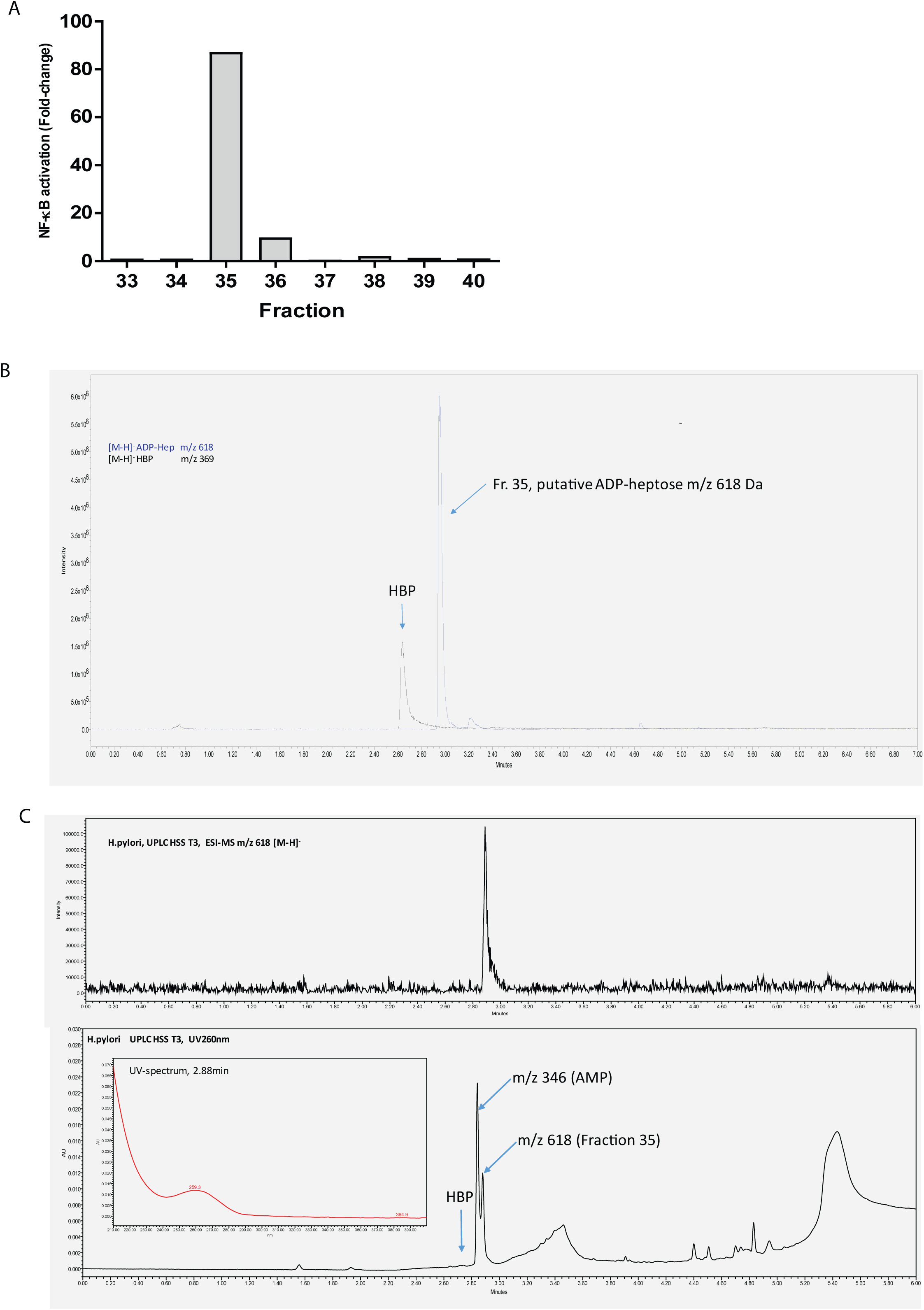
NF-κB induction capacity of specific *H. pylori* UPLC fractions and separation of *H. pylori* extract by ion-pairing reversed phase UPLC on a Waters HSS T3 column. (A) AGS NF-κB luciferase reporter cells were incubated with *H. pylori* fractions separated by ion-pairing reversed phase UPLC. Fractions were mixed with transfection reagent (lipofectamine 2000) and then added to NF-κB luciferase reporter cells for 3 h. Induction of NF-κB was determined by measurement of luciferase activity. Treatment with lipofectamine 2000 was used as negative control. Induction is shown as fold-change compared to negative control.(B) *H. pylori* extract was spiked with β-HBP. Shown are the chromatograms in SIR (Single Ion Recording) mode for m/z 369 Da and m/z 618 Da (negative mode).(C) *H. pylori* extract (carbon graphite eluate) was separated by UPLC. Shown are the chromatograms in SIR (Single Ion Recording) mode for m/z 618 Da (upper panel) and the UV absorbance at 259 nm (lower panel).

### De novo synthesis of ADP heptose

To firmly establish the identity of the 618-peak as the NF-κB-stimulating factor, we decided to synthesize β-ADP heptose both chemically and enzymatically. For chemical synthesis, we applied two strategies. We coupled β-H1P either directly to electrophilic AMP morpholidate (Wagner et al., 2009) or indirectly to activated AMP by in situ generation of 2-imidazolyl-1,3-dimethylimidazolinium chloride, which converts the 5’phosphate ester of AMP to the reactive phosphorimidazolide (Tanaka et al., 2012). Both methods worked well, yielding moderate amounts of the expected β-ADP-heptoses (L-and D-Isomers), which eluted at the same time from reversed-phase UPLC as compared to the active compound from *H. pylori* extracts.

The m/z 618 [M-H]^-^ precursor ion was further analyzed by LC-MSMS (MALDI-TOF) using chemically synthesized α-ADP-heptose as reference (Zamyatina et al., 2003). The same fragment ions could be observed in the m/z 618 precursor ion [M-H]^-^ as in the chemically synthesized α-ADP-heptose (Fig. 3A, 1 and 2, see also Supplementary Figure 3). Main fragment ions could be attributed to adenine (m/z 134), ribose-phosphate (m/z 211), heptose-phosphate (m/z 289), AMP (m/z 346) and ADP (m/z 426). MALDI-MSMS of the UPLC fractionated SPE-extract from *H. pylori* led to weak intensities of product ions (Fig 3A, 4). Alkaline phosphatase treatment of these ADP-heptoses, which were generated from the unprotected sugar phosphate, led to small amounts (1-5%) of a product with a mass of m/z 538 resulting from the loss of the heptose-phosphate group (data not shown). We presume that coupling of activated AMP to the unprotected sugar led to small amounts of an adenylylated heptose 1-phosphate product, which became esterified at the 5’-phosphate of AMP at one of the hydroxyl groups of the heptose.

**Figure 3:**
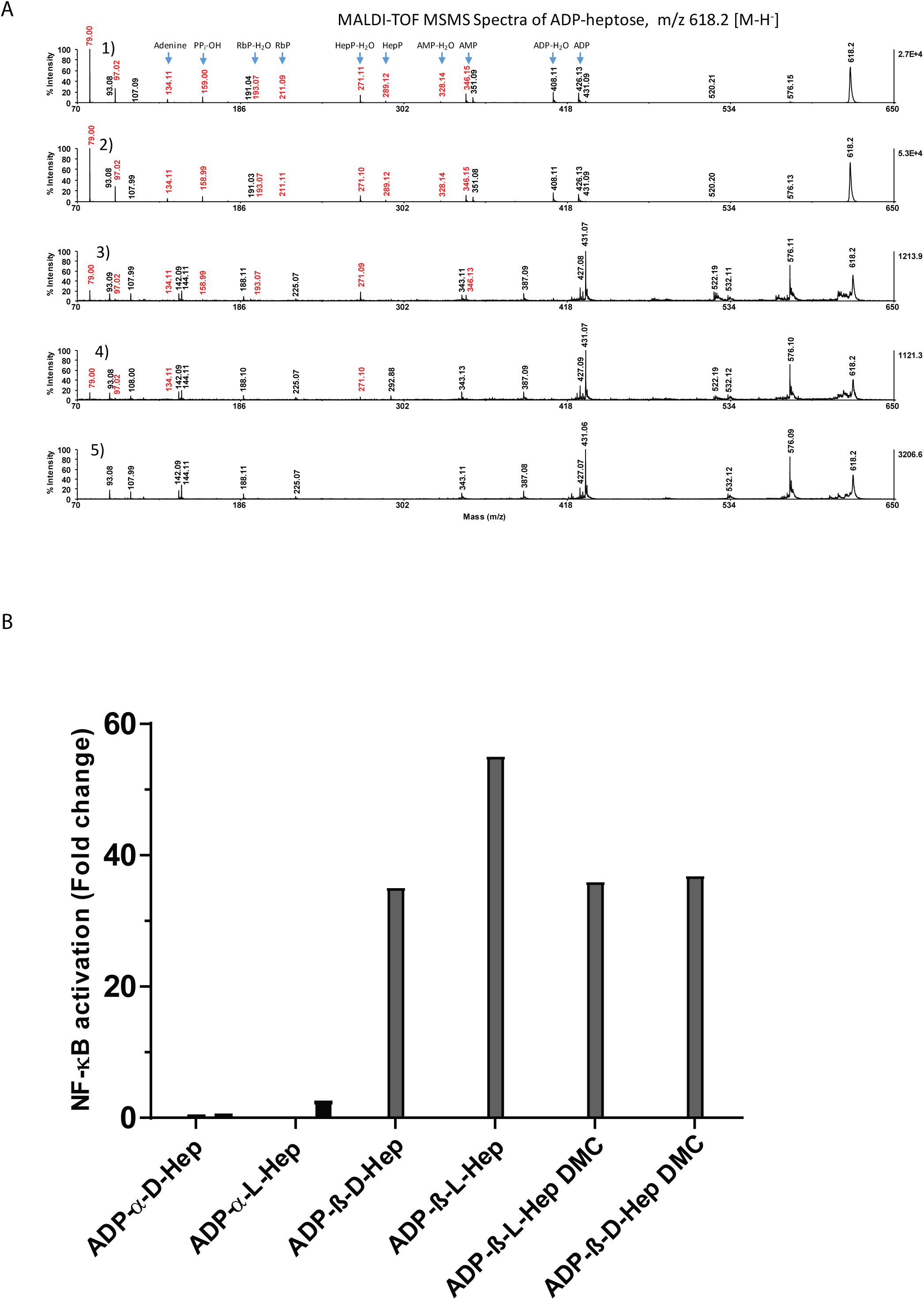
Chemically and enzymatically synthesized ADP-β-D-heptose induces NF-κB in epithelial cells. (A) MALDI-TOF MSMS spectra of ADP-heptoses, m/z 618.2 [M-H] (1) Reference spectrum: α-D-ADP-heptose (2) ADP-β-D-heptose from H1P/AMP-morpholidate synthesis (3) ADP-β-D-heptose from H1P/HldE enzymatic synthesis (4) ADP-β-D-heptose from purified *H. pylori* extract (5) Matrix control (B) NF-κB reporter cell lines were stimulated with chemically synthesized ADP-α-D-heptose and ADP-β-D-heptose, D-and L-glycero isomers, as well as with enzymatically generated ADP-β-D-heptose in D and L forms in the presence of a transfection reagent for 3 h at concentrations of 10 µM. Induction is shown as fold-change compared to untreated control. Representative data are shown.

In order to rule out the possibility that the presence of an HBP-analogue (e.g. 7’-adenylyl-heptose-1-phosphate) could be responsible for the activating effect, we also synthesized ADP-heptose, starting from the O-acetyl protected sugar (L-glycero-D-manno heptose-2,3,4,6,7-O-penta-acetate), which was phosphitylated using a phosphoramidite (see Material and Methods) and coupled by the AMP-morpholidate method as described by Zamyatina et al. (2003). Coupling of the O-acetyl protected sugar 1-phosphate to AMP-morpholidate afforded acetylated ADP-heptose (both anomeric forms), which were carefully deprotected, affording β-ADP-heptose in low yields.

As an alternative approach, we generated β-ADP-heptose enzymatically by either phosphorylating D-H7P or D/L-β-H1P with purified recombinant HldE and GmhB from *H. pylori* in the presence of ATP (Fig.3A, 3). The enzymatic conversion of the heptose monophosphates proceeded nearly quantitatively to ADP-heptose. Both enzymatically and chemically synthesized β-ADP-heptoses (D-and L-glycero isomers) were able to stimulate NF-κB, whereas α-ADP-heptose, even at higher concentrations, was not (Figure 3B).

### Pro-inflammatory activity of ADP heptose

We hypothesized that β-ADP-heptose might share the same signaling pathway as HBP, activating NF-κB via the ALPK1-TIFA axis. We therefore transfected AGS wild type as well as AGS TIFA^-/-^ and ALPK1^-/-^ knockout cells with β-ADP-heptose and measured the induction of IL-8. We observed that β-ADP-heptose-triggered NF-κB activation was completely abrogated in ALPK1 and TIFA knockout cells (Figure 4A). To further prove that β-ADP-heptose activates the ALPK1-TIFA axis, we incubated a stably tdTomato-TIFA overexpressing cell line with β-ADP-heptose. Induction of TIFAsomes could be observed in *H. pylori* wt infected cells (data not shown), cells transfected with β-ADP-heptose, and cells transfected with HBP (Figure 4B, left panel). We therefore concluded that β-ADP-heptose does indeed activate the ALPK1-triggered TIFA phosphorylation that leads to NF-κB activation.

**Figure 4:**
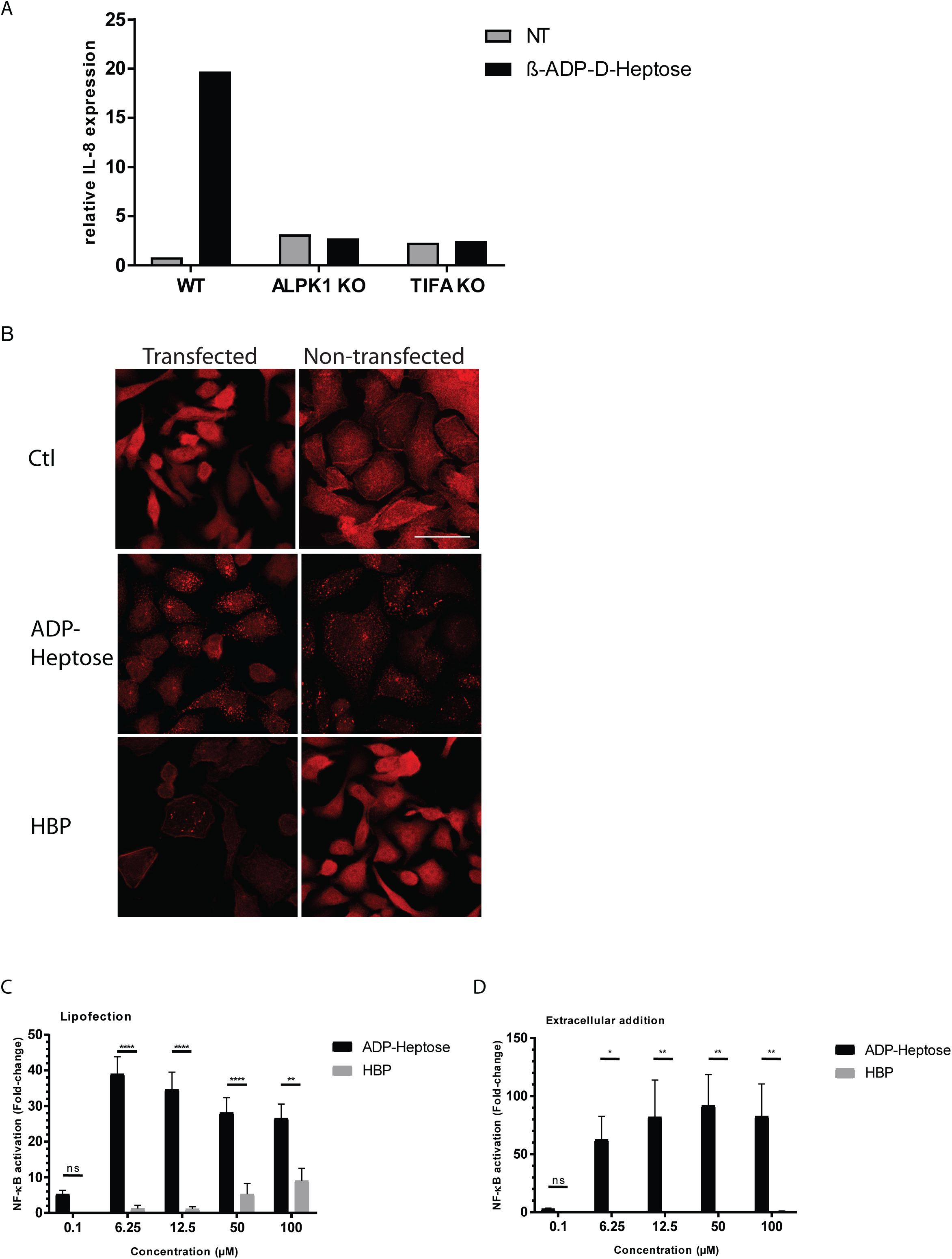
ADP-heptose-triggered NF-κB activation leads to formation of TIFAsomes and is ALPK1 and TIFA dependent. (A) ALPK1^-/-^ and TIFA^-/-^ cells were stimulated with ADP-β-D-heptose [10 µM] in the presence of a transfection reagent for 3 h or left untreated (NT) and analyzed for IL-8 induction by RT-qPCR. Representative data of at least two independent experiments are shown.(B) AGS cells stably overexpressing tdTomato-TIFA were left untreated (Ctl), treated with ADP-β-D-heptose [10 µM] or HBP [25 µM] in presence or absence of a transfection reagent for 3 h. Formation of TIFAsomes was analyzed by confocal microscopy. Scale bar: 30 µm. (C),(D) Treatment of NF-κB reporter cells with ADP-β-D-heptose and HBP in presence (C) or absence (D) of a transfection reagent for 3 h at indicated concentrations. Induction is shown as fold-change compared to an untreated control. Data represent mean ± SEM of three independent experiments.

For HBP it has been shown that delivery to the host cell cytoplasm is necessary for recognition by the host cell and subsequent NF-κB activation (Gall et al., 2017; Gaudet et al., 2015; Zimmermann et al., 2017). To our surprise, when we incubated an AGS reporter cell line with ADP-heptose in the absence of transfection reagent we observed a robust NF-κB response, whereas HBP only stimulated the cells when delivered into the cytoplasm by a transfection reagent (Figure 4C and D). This observation was confirmed in TIFA reporter cells incubated without transfection (Figure 4B right panel). Additionally, we observed that β-ADP-heptose activates NF-κB at concentrations as low as 1-10 pmol - around 100-fold lower than the concentration required for HBP (Fig. 4C and D). We conclude that β-ADP-heptose rather than HBP is the main pro-inflammatory PAMP in *H. pylori*.

## Discussion

Identifying immune stimulatory pathways is crucial for understanding how cells can sense infection and initiate an immune response. Recently, we and other groups described the ALPK1-TIFA axis stimulated by recognition of HBP as a novel pathway for the induction of an innate immune response in epithelial cells (Gall et al., 2017; Stein et al., 2017; Zimmermann et al., 2017). Yet, when analyzing *H. pylori* extracts we only detected minor amounts of HBP that appear to be insufficient for eliciting the observed NF-κB activation. Thus, we searched for additional bacterial triggers of NF-κB induction. By ion-pairing reversed phase-UPLC analysis of *H. pylori* extracts, we identified a fraction that is highly efficient in inducing an NF-κB response. Further analysis of this fraction by LC-MSMS (MALDI) revealed a prominent compound with a mass of m/z 618 [M-H]^-^ that was identified as β-ADP-heptose. The capacity of this compound to stimulate NF-κB was confirmed with chemically and enzymatically synthesized β-ADP-heptoses (D-and L-glycero isomers), both of which led to a strong NF-κB induction. Interestingly, this activation occurred even without permeabilization of the cell membranes. Activation was dependent on the presence of ALPK1 and TIFA. Hence, we show that β-ADP-heptose is present in H. pylori and acts as a novel and potent stimulator of ALPK1-TIFA controlled innate immune responses in epithelial cells.

β-ADP-heptose is a metabolite of gram-negative and also some gram-positive bacteria in the synthesis of the inner core-region of LPS (Kneidinger et al., 2002). In *H. pylori,* the *rfaE* gene encodes a bifunctional enzyme (HldE) that harbors the kinase and the ADP transferase activity responsible for generation of D-glycero-β-D-manno-heptose 1-P and ADP-D-glycero-β-D-manno-heptose (Stein et al., 2017). Previous studies have shown that deletion of *gmhA* and *rfaE* abolishes the induction of NF-κB (Gall et al., 2017; Stein et al., 2017; Zimmermann et al., 2017). Yet, a mutant in GmhB, the phosphatase converting HBP into heptose 1-phosphate, a precursor for β-ADP-heptose synthesis, still showed NF-κB activation (Gall et al., 2017; Stein et al., 2017). It is possible that this mutant is still capable of ADP-heptose synthesis via an alternative pathway, explaining the preserved NF-κB activation. Such a compensatory mechanism is known from *E. coli,* where the *gmhB* gene function can at least partially be compensated by other phosphatases (Kneidinger et al., 2002).

Former studies, including ours, have focused on HBP as the activating molecule of the ALPK1-TIFA axis, showing that permeabilization of the membrane either by lipofection or by electroporation is necessary for a robust HBP-triggered NF-κB response (Gall et al., 2017; Gaudet et al., 2015; Stein et al., 2017; Zimmermann et al., 2017). By contrast, β-ADP-heptose also activates NF-κB in epithelial cells when added extracellularly. In infection experiments, however, either a functioning type IV secretion system (Stein et al., 2017; Zimmermann et al., 2017) or cellular uptake of bacteria (Milivojevic et al., 2017) is needed to induce an immune response. This suggests that β-ADP-heptose is not normally released by bacteria but needs to be actively transported into the cell. The exact mechanism of uptake or translocation of ADP-heptose to host cells remains speculative.

Compared to HBP, β-ADP-heptose showed NF-κB activation at 100-fold lower concentrations, making this the more likely direct simulator of the ALPK1-TIFA axis. Whether HBP stimulates the same receptor only with lower affinity, or if it is converted before receptor binding, needs to be determined. Very recently, ALPK1 was identified as the direct receptor for β-ADP-heptose and shown to directly phosphorylate TIFA upon binding (Zhou et al., 2018). The same study also suggested an intracellular conversion of HBP to heptose 7-phosphate, which is also able to activate ALPK1 (Zhou et al., 2018). In support of this theory, we also observed activation in cells treated with ADP-heptose 7-phosphate (data not shown).

Here we demonstrate the presence of β-ADP-heptose in lysates of *H. pylori* and show that this compound has a substantially higher potency than HBP for inducing an NF-κB response via the ALPK1-TIFA axis. The relevance of this pathway *in vivo*, the potential usefulness of β-ADP-heptose as an adjuvant or its importance in other clinical settings are questions awaiting urgent further analysis.

## Materials and Methods

### Cell Culture

Stably tdTomato-TIFA overexpressing AGS cells (AGS JAT001), AGS cells stably expressing an NF-κB-luc2P construct (AGS JAT003), AGS SIB02 CRISPR/Cas9 control and knockout cells (AGS STZ001, AGS STZ003 and AGS STZ004) (Zimmermann et al., 2017) and the parental AGS cells were cultivated at 37 °C and 5% CO2 in RPMI (GIBCO) with 10 % heat-inactivated fetal calf serum (FCS) (Zimmermann et al., 2017).

### Bacterial Culture and Lysis Preparation

*H. pylori* WT strain P12 was grown on horse serum agar plates supplemented with vancomycin (10 µg/ml) and cultivated for two passages at 37 °C and 5% CO_2_ (Backert et al., 2000). For lysate preparation, bacteria were harvested by resuspension in water and diluted to OD_550_ 1. Cells were lysed by heating to 95 °C for 15 min. Lysates were centrifuged at 4000 x *g* for 3 min and supernatants filtered through a 0.22 µm syringe filter.

### Carbohydrate Delivery and Luciferase Assay

Stably tdTomato-TIFA overexpressing cells were starved in serum-free medium for 2 h at 37 °C and 5% CO_2_ prior to sugar delivery. Delivery of sugar into the cytoplasm was achieved by treating cells at 70% confluence with a preparation of 1 µl sugar solution mixed with 7.5 µl OptiMEM (GIBCO), 6.25 µl ATP (20 mM; Roche) and 1.25 µl lipofectamine 2000 (Thermo Fisher Scientific) after 20 min pre-incubation at room temperature (RT). Alternatively, sugars were added to the starvation medium without transfection reagent. Cells were incubated for 3 h at 37 °C and 5% CO2 and the subsequent luciferase assay performed according to the manufacturer’s instructions (Promega).

### Confocal Microscopy

Stably tdTomato-TIFA-overexpressing cells were seeded on glass coverslips and grown for 24 h at 37 °C and 5% CO2. Cells were either transfected with ADP-β-D-heptose or β-HBP using lipofectamine 2000 under serum-free conditions, treated with sugars without transfection or left untreated for 3 h at 37 °C and 5% CO2. To visualize TIFAsome formation, cells were fixated with 4% (w/v) paraformaldehyde (PFA). Coverslips were mounted with Mowiol 40-88 (Sigma, cat. No. 324590) and analyzed by laser scanning microscopy using a Leica SP8 (Zimmermann et al., 2017).

### Real Time-qPCR

RT-qPCR was performed using the Power SYBR Green RNA-to-CT 1-Step Kit according to the manufacturer’s instructions (Applied Biosystems). As described previously (Zimmermann et al., 2017), 1 ng/µl of the isolated RNA was added per reaction, the final concentration of the primers was 166 nM. RT-qPCR was performed using a StepOnePlus Real-Time PCR system (Applied Biosystems) with a one-step protocol for both RT reaction and PCR reaction. The following conditions were used during the protocol: an initial cycle for the generation of cDNA at 48 °C for 30 min; second cycle for the activation of the hot-start Taq polymerase at 95 °C for 10 min; and 40 consecutive denaturation cycles at 95 °C for 15 s, each followed by primer annealing at 60 °C for 1 min. Baseline and Cq values were determined automatically by the StepOne Software v2.3 (Applied Biosystems). Fold-changes were calculated using the 2^-δδCT^ method (Livak and Schmittgen, 2001). The following primers were used: IL-8 5′-ACACTGCGCCAACACAGAAAT-3′ and 5′-ATTGCATCTGGCAACCCTACA-3′, GAPDH 5′-GGTATCGTGGAAGGACTCATGAC-3′ and 5′-ATGCCAGTGAGCTTCCCGTTCAG-3′.

### Glycero -D-*manno*-heptoses

D-*glycero*-D-*manno-*heptose 1,7-bisphosphate (HBP, both α-and β-configuration), D-*glycero*-D-*manno*-heptose 7-phosphate (H7P), β-D-and β-L-*glycero*-D-*manno*-heptose 1-phosphate (β-H1P) were chemically synthesized as described previously (Borio et al., 2017; Guzlek et al., 2005; Zamyatina et al., 2003)

### ADP-heptose

ADP-heptoses were synthesized enzymatically with H1P or H7P as glycosyl donors, respectively, or chemically with β-H1P or with peracetylated L-*glycero*-D-*manno*-heptose. The following strategies were applied:

1. Coupling the unprotected L-and D-glycero-D-mannoheptose-1-phosphate to adenosine-5’-phosphomorpholidate (AMP-morpholidate) under anhydrous conditions in pyridine/tetrazole as described by Zamyatina et al (2003) and Wagener et. al (2009)
2. Coupling of β-H1P (D and L) to adenosine 5’-monophosphate (AMP) with 2-chloro-1,3-dichloromethylimidazolinium chloride (DMC) in D2O as described by Tanaka et al. (2012). Both, the AMP-morpholidate and the DMC/Imidazole methods were carried out in the nanomol range with up to 100 nmol H1P as starting material, keeping the reaction volume as small as possible (1-5 µl). After completion of the reaction, the mixtures were diluted with 50 µl water and ADP-heptoses were extracted with 2 vol. of chloroform/methanol (2:1). The aqueous phase was dried, the residue dissolved in 50 µl water and separated on a reversed phase HSST3 UPLC column. Products were also analyzed by LC-ESI-MS as described below.
3. L-β-ADP-heptose was synthesized according to the protocol described previously (Zamyatina et al., 2003) but with a modification regarding the introduction of the phosphate: 50 mg of the peracetylated L-*glycero*-D-*manno*-heptose were deacetylated with Hünig’s base as described. The resulting 2,3,4,6,7-penta-O-acetyl-*manno*-heptopyranose was purified by silica chromatography and phosphitylated with bis(2-cyanoethoxy)-*N,N*-diisopropylaminophosphine in dry acetonitrile/tetrazole in the presence of 1-hydroxy-6-(trifluoromethyl)benzotriazole (CF3-HOBt) (Nagata et al., 2010). The phosphitylating reagent was prepared by stirring 60 mg CF3-HOBt with 1 ml 0.1 M bis(2-cyanoethoxy)-*N,N*-diisopropylaminophosphine in anhydrous acetonitrile for 2 h under reduced pressure. The mixture was slowly added to 10 mg of the penta-O-acetyl heptopyranose in 1 ml anhydrous acetonitrile containing 3 % tetrazole over 30 min and the reaction was incubated for 8 h at RT. Oxidation to the phosphate ester was carried out by slowly adding 50 µl of 10 % tert-butylhydroperoxide over 1 h on ice. The reaction mixture was diluted with 5 vol. dichloromethane, washed with brine and 0.5 M triethylammonium bicarbonate, pH 8.5 (TEAB), successively. The organic phase was dried under nitrogen, dissolved in 500 µl acetonitrile and passed over a silica gel 60 column, which was eluted with chloroform/methanol/water (3:2:0.1), yielding 2 components with the expected molecular mass for the penta-O-acetyl-heptose-1-(bis-cyanoethyl)-phosphate of m/z 629.2 [M+Na]^+^, (ESI-MS). The later eluting sugar-phosphate (assumed to be the β-anomer) was dried and dissolved (5 mg) in 500 µl dry pyridine. Cyanoethyl groups were cleaved by adding 200 µl bis(trimethylsilyl)acetamide and 10 µl 1,8-diazabicyclo[5.4.0]undec-7-ene (DBU). The reaction was stirred for 20 min at RT (Sekine et al., 1995). The reaction was stopped by diluting with 5 vol. dichloromethane/methanol (2:1) and 2 vol. 500 mM triethylammonium bicarbonate, pH 8.2 (TEAB). The organic layer was dried, solubilized in 50% methanol and separated on an HSS T3 HPLC column that was isocratically eluted with 40% acetonitrile in 10 mM TEAB. Heptose-phosphate fractions were detected by ESI-MS (Waters QDa), dissolved in anhydrous dichloromethane, dried under vacuum over phosphorpentoxide and dissolved in anhydrous pyridine (phosphoramidite method). Coupling to AMP-morpholidate and deprotection was performed essentially as described. The resulting O-acetylated-ADP-heptose was deprotected with 50 mM triethylamine, pH 10.5 in methanol/TEAB. The product was further purified by anion exchange chromatography (Supelcosil LC-SAX 25cm x 4.6 mm; 5 µm particle). ADP-heptose was eluted with a linear gradient of 10 to 500 mM TEAB in water. Total yield: 0.65 mg (1.3%) ESI-MS (negative mode): 618, 05
4. L/D-α -ADP-heptose was synthesized as described previously (Zamyatina et al., 2003).

Synthesis of D-*glycero*-D-*manno*-heptose-7-O-ADP (D-*glycero*-D-*manno*-heptose-7-O-adenosine-5’-monophosphate ester)

The 7’-phosphate group of D-H7P was coupled to AMP via 2-chloro-1,3-dichloromethylimidazolinium chloride (DMC) in D_2_O as described by Tanaka et al (2012) as described for the synthesis of ADP-heptose from H1P.

### Enzymatic Synthesis of β-ADP-heptose from β-H1P or H7P

Up to 10 nmol D-β-H1P (triethyl ammonium salt) were incubated at 35 °C with 1 µg recombinant HldE from *H.pylori* in 20 µl HldE-Buffer (50 mM Tris HCl, pH 8.0, 150 mM NaCl, 5 mM MgCl2, 5 mM ATP) for 1 h. The reaction was stopped by addition of 50 µl chloroform/methanol (2:1). The aqueous phase was dried in a SpeedVac evaporator and dissolved in 50 µl water. The product was analyzed by LC-MS as described below. When H7P was used instead of H1P, the above reaction was performed in the presence of 1 µg of recombinant GmhB from *H. pylori*.

### Enzymatic synthesis of β-HBP from H7P

D-H7P (10 nmol, triethyl ammonium salt) was incubated with 1 µg recombinant RfaE from *N.gonorrhoeae* in 20 µl HldE-Buffer (50 mM Tris HCl, pH 8.0, 150 mM NaCl, 5 mM MgCl2, 5 mM ATP) for 1 h to 8 h at 35 °C. To destroy minor amounts of ADP-heptose, which were formed due to the presence of the endogenous *E.coli* adenylyltransferase activity, 1 µl PDE (*Crotalus adamanteus*; 3 mU/ml) was added for 30 min. The reaction was stopped by addition of 50 µl chloroform/methanol (2:1). The aqueous phase was dried in a SpeedVac evaporator and dissolved in 50 µl water. The product was analyzed by LC-MS as described below

### Derivatization of heptoses with 3-amino-9-ethylcarbazole (AEC)

Sugar-phosphates were dephosphorylated with calf intestine alkaline phosphatase (CIP) or by acid hydrolysis with trifluoroacetic acid (50 mM, 30 min, 90 °C). Reactions were conducted in 10 µl. After hydrolysis, samples were diluted with 50 µl methanol and dried in a SpeedVac evaporator. For reductive amination, the hydrolyzed samples were incubated with 100 µl AEC (60 mM in methanol) followed by the addition of 25 µl of sodium cyanoborohydride (50 mM in water) and 20 µl of acetic acid. After 1 hour at 60 °C, samples were directly submitted to UPLC-analysis. (Waters BEH C18, 50 x 2.1 mm). Compounds were eluted with a linear gradient of 10 % to 100 % acetonitrile in 0.1 % formic acid.

### MALDI-TOF MSMS

Lyophilized samples were solubilized with 4 µl 2% ACN/0.1% TFA. 1 µl of each sample was mixed with an equal volume alpha-cyano-4-hydroxycinnamic acid (0.5% solubilized in 60% ACN/0.3% TFA) and transferred onto the MALDI/MS template. MS and MSMS spectra were acquired in positive mode with a 4700 Proteomics Analyzer (AB Sciex) MALDI-TOF/TOF instrument. MS mass range was set to 300-600 Da. ADP-heptose measurements were acquired in negative mode and the MS mass range was set to 250-1000 Da. All data were evaluated manually.

### UPLC

Carbohydrates were separated by ion-pairing reversed phase chromatography on a HSS T3-UPLC column (2.1 x 150 mm, 1.8 µm, Waters) with a linear gradient of 10 mM triethylammonium bicarbonate, pH 8.5 (TEAB) from 2% acetonitrile to 90% acetonitrile (containing 10 mM TEAB) over 7 min at 45 °C and a flow rate of 0.4 ml/min. Eluted compounds were detected by UV absorbance (Waters PDA detector) and by ESI-MS detection (waters QDa) or MALDI-TOF.

The QDa was operated in an electrospray negative ion mode by applying a voltage of 0.8 kV. The cone voltage was set at 15 V. The probe temperature was set at 600 °C. A full mass spectrum between *m*/*z* 100 and 1200 was acquired at a sampling rate of 8.0 points/sec.

### Cloning and expression of the ADP-heptose-pathway enzymes RfaE, HldE, GmhB in *E.coli*

DNA recombinant procedures were performed according to standard methods described by Sambrook et al. (2001). DNA encoding the enzymes of the ADP-heptose pathway were cloned from *H. pylori* P12 (GmhB), *H. pylori* G27 (HldE) and *N. meningitidis* MS11 (RfaE). DNA templates were PCR amplified with the primers indicated below. The PCR products were digested with the indicated restriction enzymes and ligated to plasmid vector pGex-2T pre-linearized with the same restriction enzymes, yielding pGex-2T-RfaE, pGex-2T-HldE and pGex-2T-GmhB.

*E. coli* strain BL21(DE3) was used and maintained at 37 °C using Luria-Bertani (LB) medium. For strain activation, 0.5 ml of an overnight culture were inoculated into 50 ml LB medium and grown at 37 °C to an optical density of OD600 0.5. To express the GST-tagged enzymes, the three constructed recombinant plasmids pGex-2T-RfaE, pGex-2T-HldE and pGex-2T-GmhB, were transformed into *E. coli* strain BL21(DE3) to allow evaluation of protein expression. The recombinant strains were grown in LB medium in the presence of ampicillin (50 μg/ml) at 37 °C.

### Primers

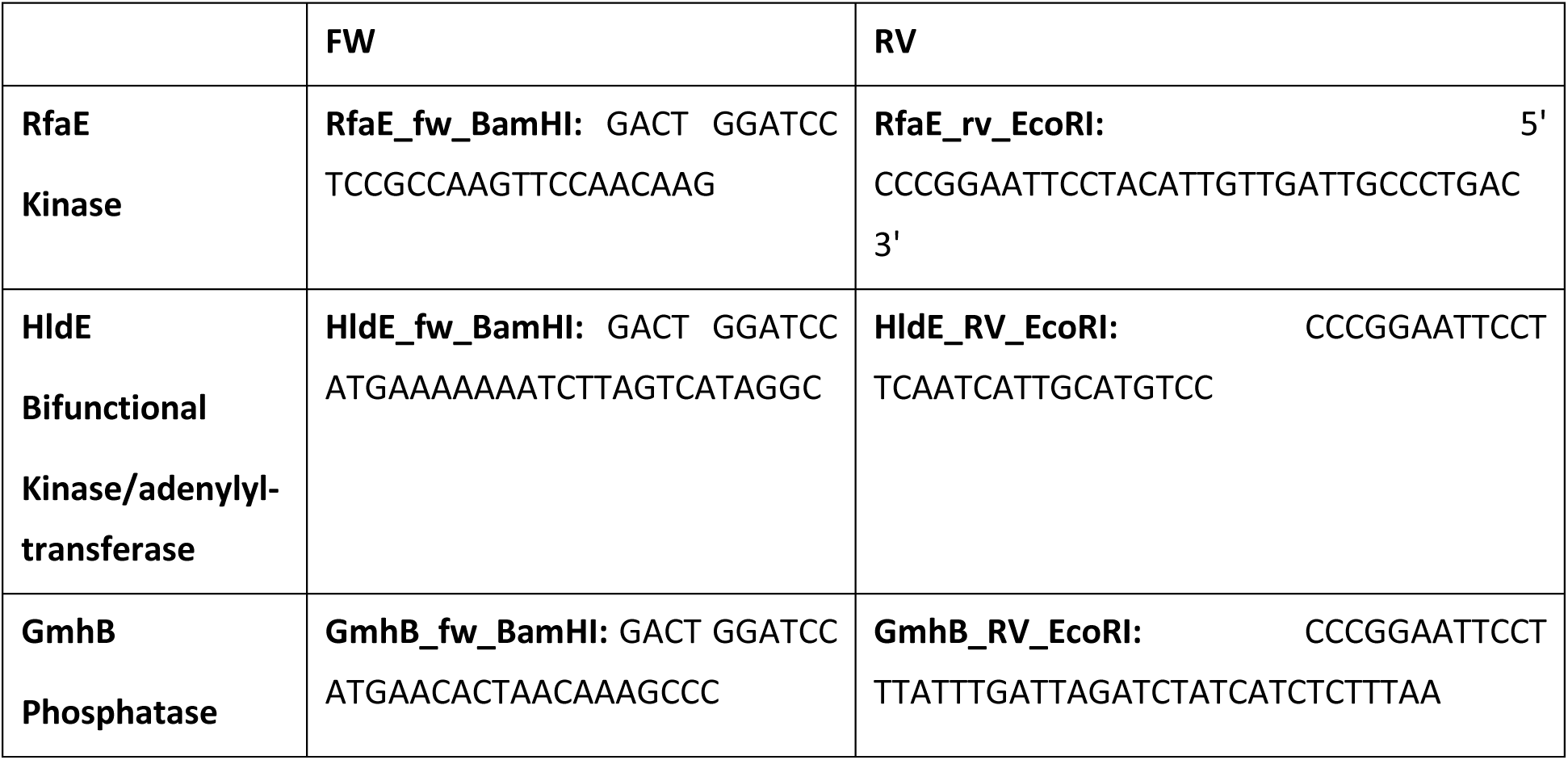

### Templates

|  |  |
| --- | --- |
| **RfaE:**<br>Acc.No AAF41238 | Strain collection: N1072 / *N. meningitidis* MS11 |
| **HldE:**<br>Acc.No O255529 | Strain collection: P224 / *H. pylori*:G27 |
| **GmhB:**<br>Acc.No D64627 | Strain collection: P511 / *H. pylori*:P12 |

#### Purification of Enzymes

Proteins were expressed in 1 l cultures, respectively. The following expression and purification protocol was applied per 100 ml of bacterial culture: 5 ml pre-culture was grown for 16 h at 37 °C, followed by the addition of 95 ml of pre-warmed LB medium with ampicillin. At the optical density (OD) of 1.5 at 600 nm, protein expression was induced for 4 h at 30 °C with 1 mM IPTG. Cells were harvested by centrifugation (4000 × *g*, 4°C, 15 min). The cell pellet was frozen in liquid nitrogen thawed on ice for 15 min and resuspended in 3 ml of lysis buffer (20 mM Tris, pH 8.0, 5 mM EDTA). 50 μl of lysozyme solution (10 mg/ml) was added and incubated on ice for 30 min. 1 μl of benzonase grade II (Merck,1 U/µl) and 50 µl of 1 M MgCl2 was added and incubated on ice for 30 min. The lysates were centrifuged at 6000 × g, 4 °C, 30 min. The supernatants were mixed with 200 µl of glutathione-sepharose 4B beads (50 % suspension in PBS) for 4 h, washed several times with PBS. Bound proteins were eluted with 20 mM glutathione in 100 mM Tris HCl, pH 8.0. Eluates were applied on a sephadex G20 column equilibrated in PBS to get rid of glutathione. Protein concentration was adjusted to 10 mg/ml by ultrafiltration centrifugal devices and stored frozen in aliquots at −80 °C.

### Isolation of ADP-heptose from *H. pylori*

*H. pylori* strain P12 was grown on horse serum agar plates supplemented with vancomycin (10 µg/ml) and cultivated for two passages at 37 °C and 5% CO2 (Backert et al., 2000). For lysate preparation, bacteria were harvested by resuspension in water and diluted to OD_550_ 1. Cells were lysed by heating to 95 °C for 15 min. Lysates were centrifuged at 4000 x *g* for 3 min and supernatants filtered through a 0.22 µm syringe filter.

One ml of the cleared lysate was extracted by the addition of 2 ml chloroform/methanol (2:1). After vortexing and centrifugation (5 min, 10,000 x *g*), the aqueous phase was passed over a HyperSep SPE-Aminopropyl cartridge (200 mg/3 ml) which was equilibrated with 50 mM acetic acid in 50% methanol. Bound compounds were eluted with 500 mM TEAB, pH 8.5 in 50% methanol. Eluates were dried in a SpeedVac evaporator, solubilized in 1 ml 10 mM ammonium bicarbonate, pH 8.0 and passed over a graphite carbon column (Supelclean ENVI-Carb, 1 ml). Sugars were eluted with 30% acetonitrile. The eluate was dried in a SpeedVac evaporator, dissolved in 10 µl 10 mM TEAB-buffer and loaded onto a HSS-T3 reversed phase UPLC column (3×150 mm, 1.8 µm) equipped with a HSS T3 VanGuard Precolumn. Compounds were separated at a flow rate of 0.4 ml/min at 45 °C with a linear gradient from 2% acetonitrile to 80% acetonitrile containing 10 mM TEAB as ion-pairing reagent. Fractions were analyzed by UV (PDA-Detector).

## Acknowledgements

The authors thank Alla Zamyatina for providing samples of ADP α-heptoses, Meike Soerensen for technical assistance and Rike Zietlow for editing the manuscript.

